# Highly dynamic supernumerary mini-chromosomes in a *Magnaporthe oryzae* strain

**DOI:** 10.1101/2024.12.10.627822

**Authors:** Guifang Lin, Huakun Zheng, Dal-Hoe Koo, Zonghua Wang, David Cook, Barbara Valent, Sanzhen Liu

## Abstract

*Magnaporthe oryzae* (syn. *Pyricularia oryzae*), the causative agent of devastating crop diseases, exhibits remarkable genomic plasticity that contributes to its adaptability and pathogenicity. Individual *M*. *oryzae* strains may contain supernumerary mini-chromosomes, which are dispensable and highly repetitive. Here, we explored the stability of two mini-chromosomes of a *Lolium* strain isolated in the US, TF05-1, in which one mini-chromosome contains sequences nearly identical to the mini-chromosome of the wheat isolate B71 from Bolivia. The discordance of their phylogenetic relationships based on genomic polymorphisms in core chromosomes and polymorphisms in mini-chromosomes indicated horizontal transfer of the mini-chromosome. Karyotyping analysis and genome sequencing analysis found variation in numbers and sizes of mini-chromosomes among asexual monoconidial progeny of TF05-1. Rearrangement within mini-chromosomes occurred frequently in the TF05-1 progeny. We characterized an intrachromosomal rearrangement presumably mediated by a palindrome repeat. The rearrangement resulted in a 300-kb deletion and a 900-kb duplication. The susceptibility to structural variation in mini-chromosomes may be associated with repetitive features and the high activity of transposable elements in mini-chromosomes, in which many intact retrotransposons were recently inserted, largely unmethylated, and likely have yet to be silenced by fungal genome defense mechanisms such as repeat-induced point mutation.

## Introduction

Genome plasticity mediated by genome rearrangement and the horizontal transfer of supernumerary chromosomes constitutes an important mechanism driving the adaptive evolution of fungal pathogens [1–5]. Supernumerary chromosomes, also known as accessory, dispensable, or B chromosomes, are extra chromosomes that exist in some but not all individuals of a species [2]. These extra chromosomes are rich in repetitive sequences, which are largely transposable elements that facilitate genome rearrangement [6,7]. Supernumerary chromosomes are often smaller than indispensable core chromosomes [8]. The relatively small size may facilitate horizontal transfer through heterokaryosis [9]. Although supernumerary chromosomes are documented to be highly dynamic under meiosis in sexual cycles [8,10–12], the stability of this repetitive genome compartment during vegetative growth is less clear.

*Magnaporthe oryzae* (syn. of *Pyricularia oryzae*) is an agriculturally important species complex that serves as an ideal model system to study genomic dynamics associated with supernumerary chromosomes. *M. oryzae* infects a wide range of grasses, causing destructive blast diseases. *M. oryzae* isolates are grouped into various lineages, known as host-adapted pathotypes, according to the primary host plant genera they infect. The *Oryza* pathotype (MoO) on rice, the *Setaria* pathotype (MoS) on foxtail millet, and the *Eleusine* pathotype (MoE) on finger millet have long threatened food production, whereas the *Triticum* pathotype (MoT) on wheat and the *Lolium* pathotype (MoL) on ryegrass emerged as agricultural threats within the last 50 years [13–15]. In *M*. *oryzae*, supernumerary chromosomes range from hundreds of kilobases to approximately 3 Mb in size. They are referred to as mini-chromosomes because indispensable core chromosomes are typically larger than 3 Mb [8]. Similar to supernumerary chromosomes in other fungal species, mini-chromosomes in *M. oryzae* are enriched in repetitive sequences, but they also contain genes. Indeed, *in planta* specifically expressed effector genes have been identified in mini-chromosomes [16]. Analysis through contour-clamped homogeneous electric field (CHEF) electrophoresis, cytological approaches, or prediction approaches using whole genome sequencing (WGS) data revealed the prevalence of mini-chromosomes in isolates from various *M*. *oryzae* pathotypes [8,10,16–20]. Such mini-chromosomes contribute to genomic dynamics in *M*. *oryzae*. Mini-chromosomes in *M*. *oryzae* strains have been found to exhibit non-Mendelian inheritance in genetic crosses and they were prone to genome rearrangement [8,10]. Repetitive sequences are attributed to creating genomic environments conducive to rapid genome evolution by offering homology for duplication, loss, and genome rearrangement [21]. Clonal strains from the pandemic B71 phylogenetic branch causing wheat blast outbreaks across continents exhibited significant variation in the number of mini-chromosomes and their DNA content [22]. In addition, mini-chromosomes appear to be capable of exchanging DNA with core chromosomes, reshuffling effector genes and repetitive sequences [16–18]. To further explore stability of mini-chromosomes and their contribution to genome dynamics during vegetative growth by *M. oryzae*, we identified a native U.S. *Lolium* isolate that contained a mini-chromosome closely related to the B71 mini-chromosome. We developed and applied Fluorescence In Situ Hybridization (FISH), a technique allowing for the detection of specific DNA sequences on chromosomes [23]. Application of FISH together with CHEF, an experimental evolution assay, and genomic approaches has documented remarkable dynamics of mini-chromosomes in a U.S. MoL isolate causing the recently emerged gray leaf spot disease on turf and forage grasses.

## Results

### *Lolium-*adapted isolate TF05-1 shared highly similar mini-chromosome sequences to wheat-adapted isolate B71

Whole genome sequencing (WGS) Illumina short-read data of MoT isolates (N=13), MoL isolates (N=6), and a MoE isolate were processed to identify SNPs using the MoT B71 genome as the reference (**Table S1**) [16]. In total, 192,227 biallelic SNPs were found among these isolates. Note that all these isolates were previously predicted to carry at least one mini-chromosome [20]. We first extracted 184,838 SNPs on core chromosomes, from chromosome 1 to 7, to construct a phylogenetic tree (**Figure 1A**). In the tree, MoT and MoL isolates were clustered separately into two clades. SNPs (N=7,389) on the mini-chromosome were used to construct another tree (**Figure 1B**), from which MoT and MoL isolates were not completely separated. Notably, MoT strain B71, isolated in Bolivia in 2012, was clustered together with MoL strain TF05-1, which was isolated in Kentucky US in 2005, with the bootstrap confidence level of 100. On core chromosomes, 51.3% SNPs had different genotypes between TF05-1 and B71. In contrast, on the mini-chromosome, only 0.2% SNPs showed distinct genotypes between the two isolates (**Table S2**). Phylogenetic and SNP analyses indicated that the TF05-1 genome contained genomic sequences that were highly similar to the B71 mini-chromosome, compared to more divergent sequences observed on core chromosomes.

**Figure 1.**
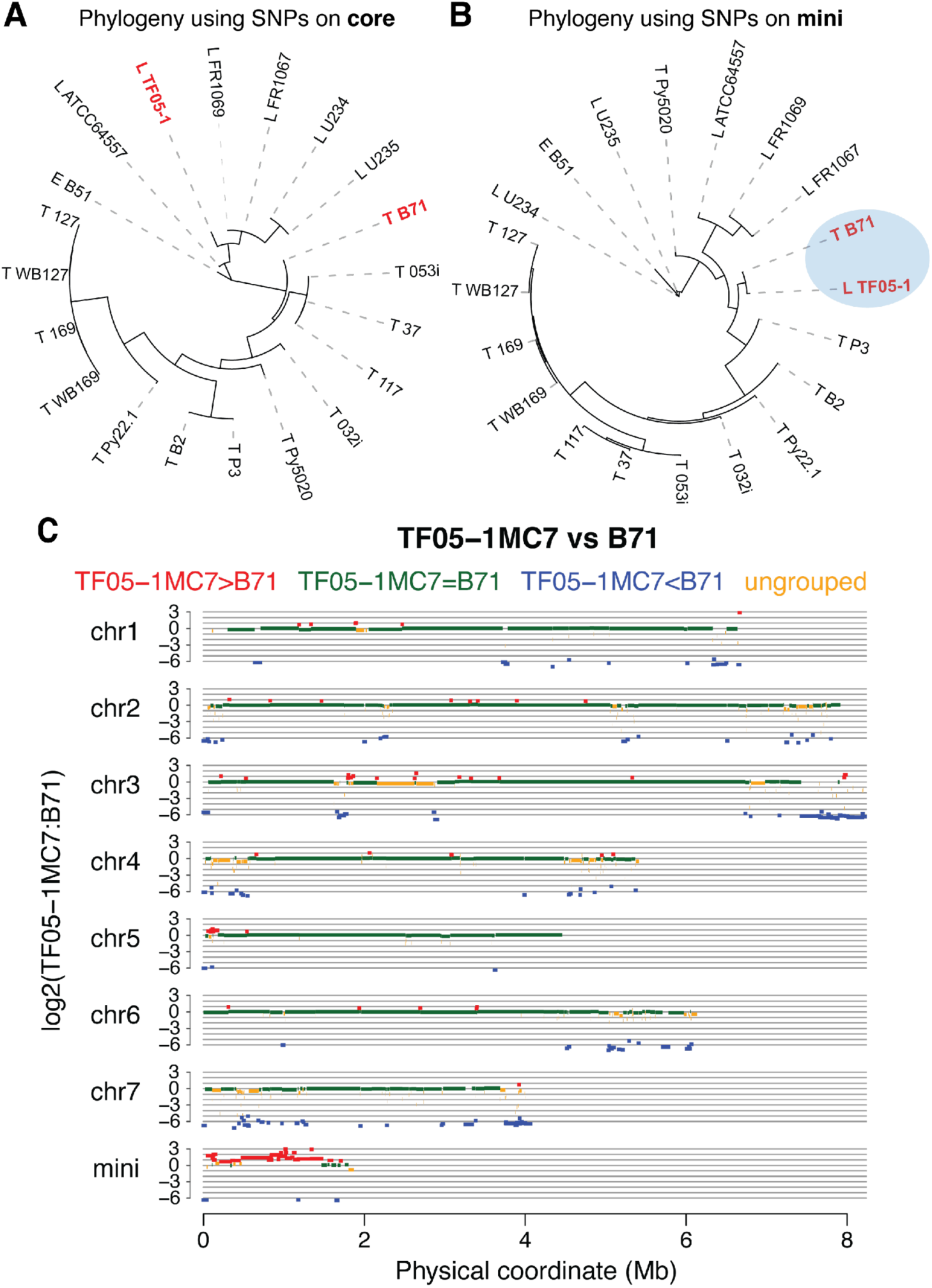
TF05-1 contains highly similar sequences to the B71 mini-chromosome. (**A**, **B**). Nineteen Mo strains isolated from either *Triticum* (T) or *Lolium* (L) and a strain B51 from *Eleusine* (E) were used to construct two phylogenetic trees. The B71 genome, including a mini-chromosome, served as the reference genome. (**A**). SNPs on core-chromosomes were the input for the tree construction. (**B**). SNPs on the mini-chromosome were the input. (**C**). Copy number comparison between TF05-1MC7 (MC7) and B71 using Illumina short reads through the CGRD pipeline. Red, green, blue segments indicate higher, equal, lower copies in TF05-1MC7 as compared to B71. Orange segments represent undetermined genomic regions.

WGS Illumina data of TF05-1 and B71 were used to examine copy number variation between the two genomes using the Comparative Genomic Read Depth (CGRD) pipeline (**Figure 1C**, **Tables S3**, **S4**) [24]. On core chromosomes, 131 regions spanning 2.2 Mb, which accounts for 5.7% of B71 core chromosomes, were found to be absent in TF05-1. The large absent regions included the ends of chromosomes 3 and 7. Three small regions, comprising 4.9% of the B71 mini-chromosome, were absent in TF05-1. In addition, CGRD analysis showed that most regions of the B71 mini-chromosome were duplicated in TF05-1, indicating that TF05-1 might contain multiple mini-chromosomes.

### TF05-1 contained mini-chromosomes that exhibited high levels of instability

Two monoconidial isolates, MC5 and MC7, were derived from TF05-1 and karyotyped through CHEF electrophoresis to assess the presence of mini-chromosomes (**Figure 2A**). The CHEF gel showed MC5 and MC7 appeared to share a small mini-chromosome with the size between 1.12 Mb and 1.6 Mb. MC5 contained an additional larger chromosome of the size larger than 2.2 Mb, which was not observed in MC7.

**Figure 2.**
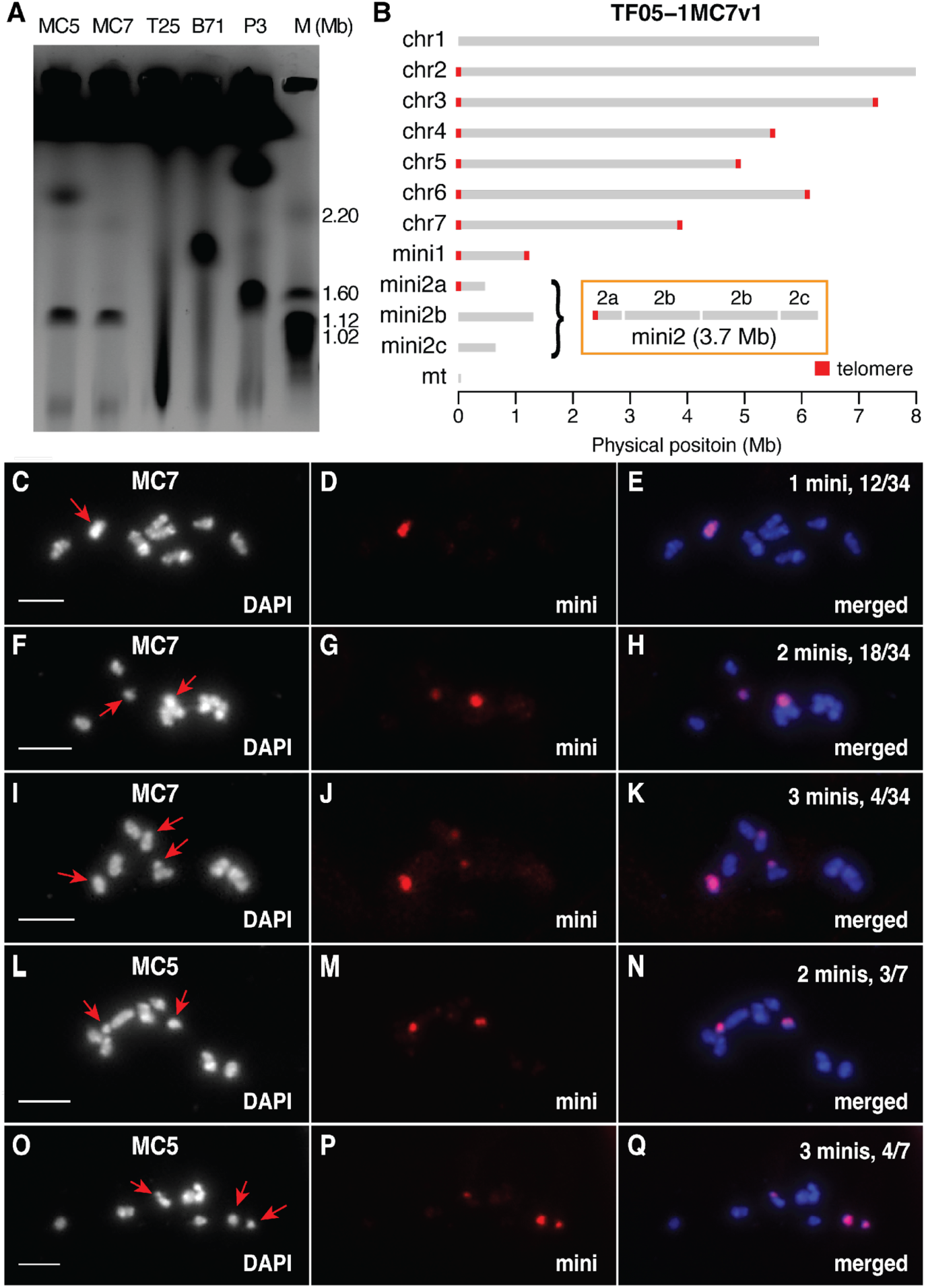
Mini-chromosomes in MC7 evidenced with CHEF, genome assembly, and FISH. (**A**). CHEF gel of TF05-1MC5 (MC5), TF05-1MC7 (MC7), T25, B71, and P3. Note that T25 had no mini-chromosomes, B71 and P3 possessed one and two mini-chromosomes, respectively. (**B**). The contig view of the near-finished genome assembly of TF05-1MC7. Based on the assembly graph and supporting sequencing depths, three contigs (mini2a, mini2b, and mini2c) were hypothesized to be from the large mini-chromosome (mini2) and mini2b was duplicated, which is illustrated in the orange box. (**C-H**) Identification of mini-chromosomes in MC7 (**C-K**) and MC5 (**L-Q**) by fluorescence in situ hybridization (FISH) using mini-chromosome probes (mini). The number of mini-chromosomes (minis), indicated at the top right of the merged images, was inferred for each observed nucleus. Red arrows in the DAPI images on the left indicate mini-chromosomes. The number of germ tube cells with the same number of mini-chromosomes inferred is listed with the total cells observed for an isolate (e.g., “2 minis, 18/34” in **H** stands for 18 out of 34 cells containing two minis). Bars=5 µm.

Nanopore long reads were produced for genome assemblies of both monoconidial isolates. Illumina WGS data were used for polishing the assembled genomes. The MC7 genome assembly (MC7v1) resulted in DNA sequences of 45.5 Mb in total, consisting of seven core chromosomes, a telomere-to-telomere contig of 1.2 Mb, a circular mitochondrial genome, and three additional contigs (**Figure 2B**, **Table S5**). We previously developed an approach to analyze gel bands of the small chromosome through excising, purifying, and sequencing with an Illumina sequencing platform [16]. Here, the approach was further combined with the CGRD pipeline for associating CHEF DNA bands with assembled contigs. Analysis of the one observed MC7 mini-chromosome and both MC5 mini-chromosomes using the MC7 genome assembly showed that both the visible MC7 mini-chromosome and the smaller MC5 mini-chromosome were associated with the 1.2-Mb contig with telomeres at both ends (**Figure S1A, S1B**). This contig was therefore referred to as mini1. Three MC7 contigs were associated with the larger mini-chromosome of MC5, suggesting that MC7 contained mini-chromosome sequences in addition to mini1 (**Figure S1C**). These three MC7 contigs were named mini2a, mini2b, and mini2c. Read depth analysis showed that mini2b was assembled with approximately twice the read depth as other chromosomal contigs, indicating that mini2b was duplicated in MC7 (**Table S5**). Collectively, we hypothesized that MC7 contained an additional mini-chromosome of 3.7 Mb in size (mini2), consisting of mini2a, duplicated mini2b, and mini2c (**Figure 2B**). The large size of the mini-chromosome explained why it could not be well separated from core chromosomes by CHEF.

Nanopore long read sequencing and genome assembly were also conducted for MC5. Assemblies of core chromosomes of MC5 were nearly complete, with 13 of 14 chromosomal ends containing telomere repeats (**Figure S2**, **Table S6**). In contrast, mini-chromosomes were highly fragmented. Mini1 sequences were found in five contigs, and 13 contigs might account for other mini-chromosome sequences. The genome assembly of MC5 did not fully resolve the complexity of the MC5 genome.

CGRD analysis comparing MC5 with MC7 did not find copy number changes in core chromosomes or mini1 of MC7. The analysis indicated copy number gain in MC7 mini2 (**Figure S1D**). Sequencing analysis of the DNA recovered from the visible MC5 mini-chromosome band larger than 2.2 Mb showed that it contains most sequences of MC7 mini2 (**Figure S1C**). Collectively, our data indicated the MC5 genome harbored MC7 mini1 and at least one or two additional mini-chromosomes, which were likely the rearrangement products of mini2 of MC7.

### FISH analyses identified mini-chromosome variations in MC5 and MC7 germ tube cells

To further understand distinct karyotypes of MC5 and MC7, we employed cytological karyotyping using the germ tube burst method (Ayukawa and Taga, 2022) and incorporated FISH, which allows for the analysis of chromosome numbers and size within individual fungal cells. Probes for mini-chromosomes were prepared from DNAs recovered from the mini-chromosome bands in CHEF gels. Chromosome painting, using mini-chromosome DNA as probes, surprisingly revealed dynamic numbers of mini-chromosomes in different conidial germ tube cells from both MC7 and MC5 cultures (**Figures 2C-2Q**). Among the 34 MC7 nuclei observed, 12 (35.3%) contained 1 mini-chromosome (**Figures 2C-2E**), 18 (52.9%) contained 2 mini-chromosomes (**Figures 2F-2H**), and 4 (11.8%) contained 3 mini-chromosomes (**Figure 2I-2K**). A similar variation was observed in the MC5 genome, where 3 nuclei (42.9%) contained 2 mini-chromosomes (**Figure 2L-2N**), while 4 nuclei (57.1%) contained 3 mini-chromosomes (**Figures 2O-2Q**). Chromosome-specific probes were developed for each core chromosome (**Table S7**). FISH with probes derived from the core chromosome showed no variation in core chromosome numbers across all the cells observed for MC5 and MC7 (N=20) (**Figure S3**).

### B71 mini-chromosome sequences were largely found in MC7, despite rearrangements

The MC7 genome was compared with the B71 reference genome through NUCMER alignments followed by SYRI for discovery of the synteny and structural variation [24,25]. Genome comparison showed that synteny was well maintained between core chromosomes in MC7 and B71. Within syntenic blocks, three large insertions and one large deletion in MC7 were identified, including an insertion of 482 kb on chromosome 5 (**Figure 3**, **Table S8**). We found no other major structural variation events except two small inversions on core chromosomes (**Figure 3A-G**). Only 1% of MC7 core chromosome sequences could not be aligned to B71 core chromosomes. Most of the unaligned regions were located at the ends of core chromosomes. Large regions on the ends of chromosomes 3 and 7 of B71 were not aligned with MC7, consistent with the findings from the CGRD comparison.

**Figure 3.**
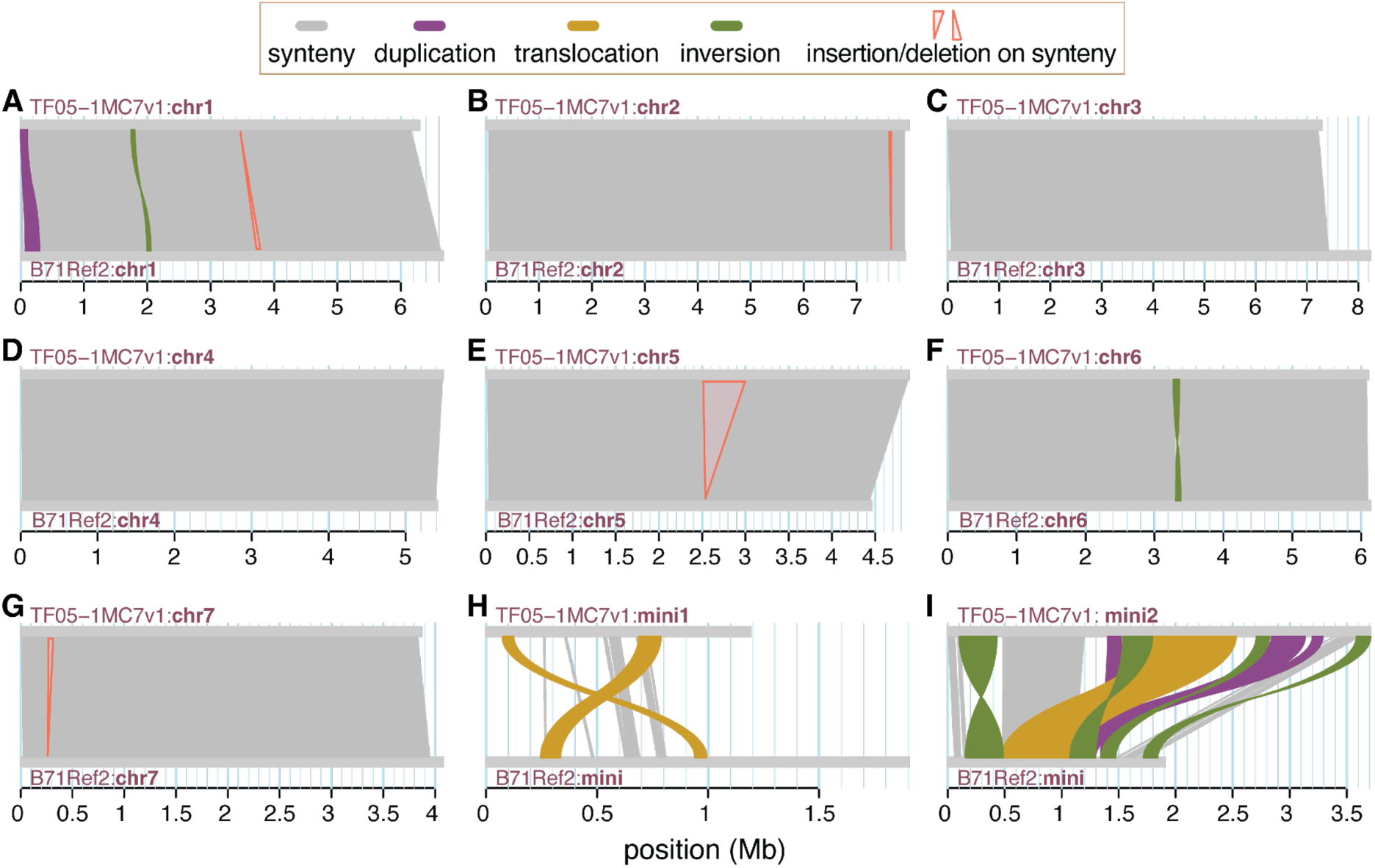
Synteny and structural variation between TF05-1 and B71. (**A**-**I**) Chromosomal comparisons of pairs from the TF05-1MC7 gnome assembly and the B71 reference genome were plotted. Genomic alignments were performed with NUCMER followed by Syri analysis to identify synteny (gray) and structural variation with at least 50 kb, including inversion (green), translocation (orange), and duplication (purple). The duplication at the end of chromosome 1 was likely to be the incomplete assembly of ribosome DNA repeats. Insertion and deletion events larger than 30 kb in syntenic regions were labeled in red.

In contrast to the syntenic conservation of core chromosomes, MC7 mini-chromosomes appeared to be structurally divergent. MC7 mini1 was highly divergent from the B71 mini-chromosome with 57.5% of sequences unaligned, while 11.2% of MC7 mini2 sequences could not be aligned to the B71 mini-chromosome (**Figure 3H, 3I**, **Table S9**). As mini2 of MC7 included a duplication of mini2b, duplicated and translocated regions were expectedly identified on mini2 in comparison to the B71 mini-chromosome. Both MC7 mini-chromosomes carried the genomic segment containing two effector genes, *BAS1* and *PWL2*, on syntenic regions (**Figure S4**). The *BAS1*-*PWL2* segment on mini1 included an insertion between the two genes. The insertion was not present in the *BAS1* and *PWL2* region in either mini2 or the B71 mini-chromosome. SNPs between MC7 and B71 were identified in syntenic regions. Mini1 syntenic regions accumulated a higher density of SNPs, 5.7 SNPs per kb, than syntenic regions on core chromosomes whose SNP density was 2.8 SNPs per kb. On the contrary, mini2 regions syntenic with the B71 mini-chromosome accumulated fewer SNPs at the level of 0.5 SNPs per kb. The results suggested MC7 mini2 was genetically closer to the B71 mini-chromosome than either the MC7 mini1 or core chromosomes.

### Frequent intra-mini-chromosomal rearrangements were observed in the MC7 progeny

Monoconidial isolate MC7 was further grown to fill two oatmeal agar plates (defined as two generations), followed by single spore isolation (**Figure 4A**). In total, cultures grown from 40 second-generation single spores were analyzed by CHEF karyotyping. Two mutant isolates MC7202 and MC7212 displayed different mini-chromosome karyotypes compared to the original MC7 (**Figure 4B**). MC7202 had a mini-chromosome larger than the 1.2 Mb mini1 in MC7. MC7212 contained a mini-chromosome sized between 2.4 Mb and 2.7 Mb in addition to a smaller one similar to mini1. FISH examination further indicated a high level of instability of mini-chromosomes in both MC7202 and MC7212. Of MC7202, two mini-chromosomes were observed in 18 nuclei (69.2%), and one mini-chromosome was observed in the remaining 8 (30.7%) nuclei (**Figure 4C-H**). Of MC7212, cells containing zero, 1, and 2 were observed in 1 (8%), 9 (69.2%), and 3 (23.0%) nuclei, respectively (**Figure 4I-Q**).

**Figure 4.**
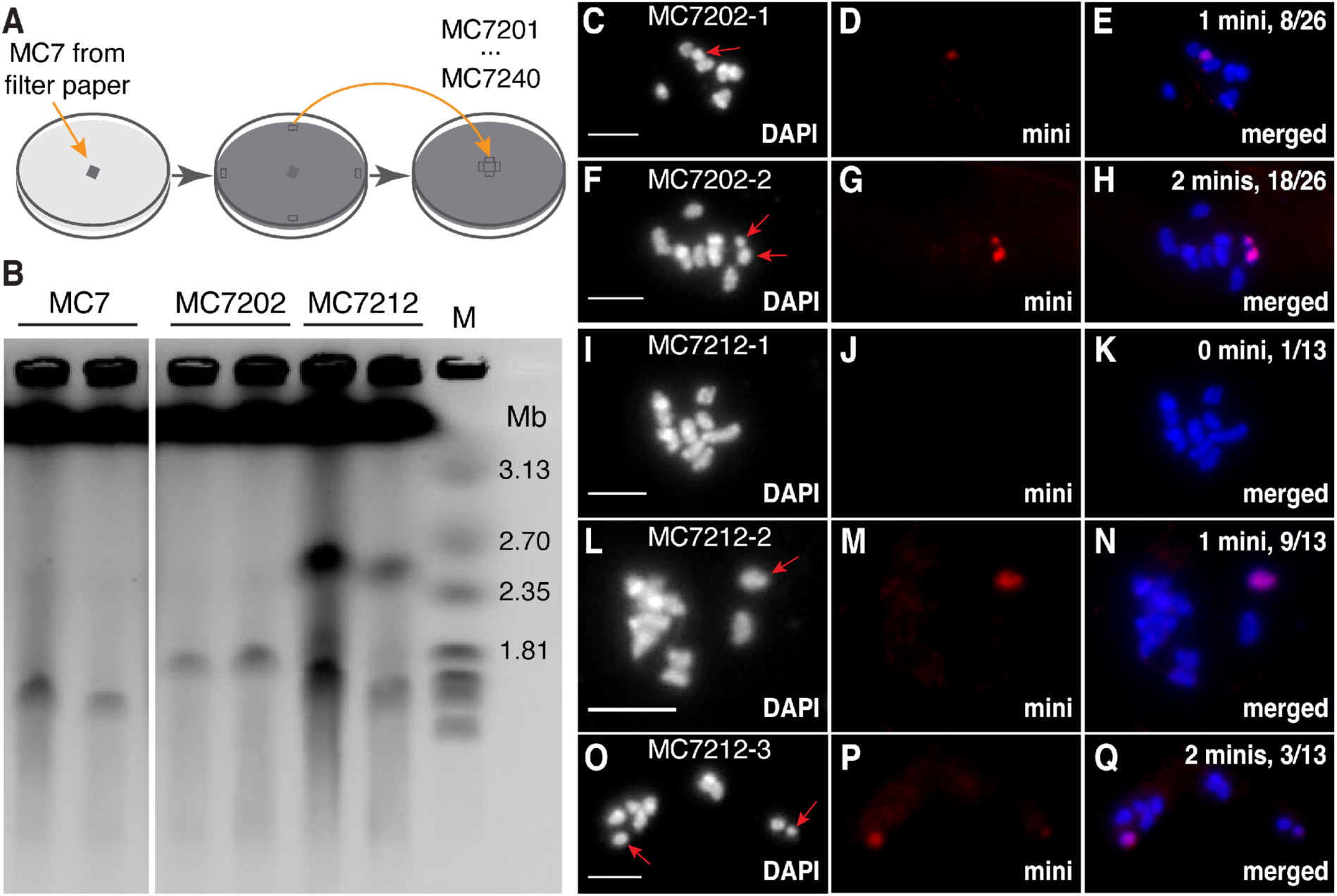
Variation of mini-chromosomes in MC7 progenies. (**A**). Purified TF05-1MC7 was cultured on oatmeal agar plates and plugs of hyphae from the edges of the first plates were transferred to new “second generation” plates as shown. Forty single spores were isolated from the edges of these second plates and labeled MC7201 through MC7240 to indicate the growth generation and spore number. (**B**). CHEF gel results of MC7; and from MC7 progeny strains MC7202 and MC7212. (**C-Q**). FISH results of MC7202 and MC7212. Chromosome painting using mini-chromosome DNAs as probes (mini) revealed variation in the number of mini-chromosomes among MC7202 and MC7212 conidial germ tube cells. Germ tubes from MC7202 contained either one (MC7202-1) or two (MC7202-2) minis and germ tubes from MC7212 contained one (MC7212-1), two (MC7212-2), or three (MC7212-3) minis. Quantification of the number of inferred mini-chromosomes (minis) is indicated at the top right of the merged images (e.g., “2 minis, 3/13” in **Q** stands for 3 out of 13 total cells examined containing two minis). Red arrows indicate mini-chromosomes in the DAPI images on the left. Bars=5 µm.

Nanopore long reads were generated for both MC7202 and MC7212. We first adapted CGRD using long reads as the input for analyzing copy number variation. MC7202 CGRD showed a >300 kb deletion from the beginning of MC7 mini1, and a duplication of the rest of mini1 (**Figure 5A**). Consistently, the deletion region of mini1 was not in the MC7202 genome assembly (MC7202v1) (**Figure S5**, **Table S10**). The duplication region detected was assembled into at least two contigs and their overall read depth was approximately twice as high as the depths for core chromosomes. We found a 3.2-kb tandem inverted repeat near the rearrangement site on MC7 mini1, forming a 6.6-kb palindrome structure (**Figure 5B**). Based on a known model for repairing DNA double stranded breaks near a palindrome, the duplication was hypothesized to be the product of intrastrand base pairing of the tandem inverted repeat (**Figure 5C**) [26].

**Figure 5.**
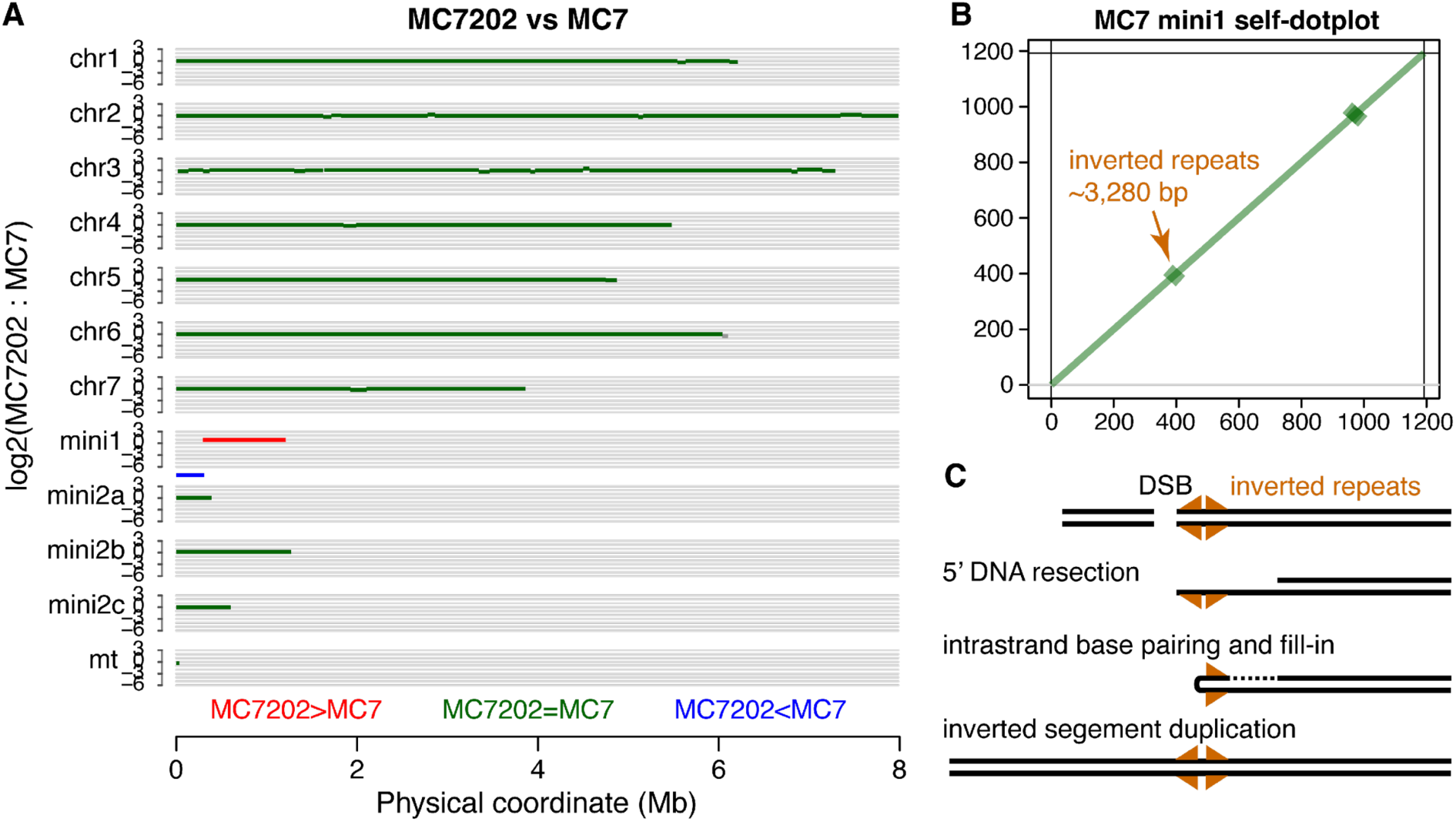
Structural changes in mini1 of TF05-1MC7202. (**A**). CGRD comparison between isolate TF05-1MC7202 (MC7202) and the parental isolate TF05-1MC7 (MC7). The comparison found MC7202 mini1 contained a large deletion from 1 bp to 309 kb as well as a large duplication from 309 kb to 1.19 Mb. (**B**). A self-comparison dotplot of mini1 in TF05-1MC7, which identified two inverted repeats, from 390 kb to 397 kb. (**C**). A model explains how the structural alteration occurred in mini1 of MC7202.

MC7212 CGRD compared with MC7 did not find drastic copy number variation. The MC7212 genome assembly (MC7212v1) showed that it contained a mini-chromosome that was well assembled with telomere repeats on both ends and it was nearly identical to MC7 mini1 (**Figure S6A**, **Table S11**). The length of mini1 was consistent with the size of the small chromosome estimated from the CHEF result. The large mini-chromosome in MC7212 was expected to be smaller than mini2 in MC7 based on CHEF. However, sequences from mini2a, mini2b, and mini2c of MC7v1 were all nearly covered in the MC7212v1 assembly (**Figure S6B**). Read depths for the assembled contigs indicated that MC7212 maintained a partial duplication of mini2b and acquired duplications of partial sequences of mini2c (**Table S11**). Therefore, partial duplications and deletions were hypothesized to result in a smaller mini2 in MC7212, which was visible in CHEF karyotyping (**Figure 4B**).

### Most LTR retrotransposons on mini-chromosomes were recently inserted and hypomethylated

Annotation of transposable elements revealed 45 intact long terminal repeat (LTR) retrotransposons categorized to *Gypsy* (N=22), *Copia* (N=11) and unknown (N=12) superfamilies (**Table S12**). Of them, 40% were located on mini-chromosomes comprising 8% of the genome. Insertion time of LTR retrotransposons can be estimated by identifying polymorphisms between two LTRs at ends of a retrotransposon because identical LTRs are produced during the insertion event [27]. Insertion times ranging from 0 to 3.3 million years were estimated for the LTR retrotransposons. Insertions of the 45 LTR retrotransposons were grouped into old or young if the insertion time was older or younger than 200,000 years, respectively. By using this cutoff, at most one polymorphic base pair between the two LTRs was allowed for a young LTR insertion. Old and young LTR insertions were not proportionally distributed on core chromosomes or mini-chromosomes (χ^2^=4.15, *p*-value=0.042), of which only 2/14 insertions on mini-chromosomes were old. Interestingly, the insertion time was correlated with G+C content (**Figure 6A**). Most young LTR retrotransposons maintained G+C content at levels higher than 50%. On the contrary, G+C content of almost all LTR retrotransposons older than 1 million years were lower than 46%.

**Figure 6.**
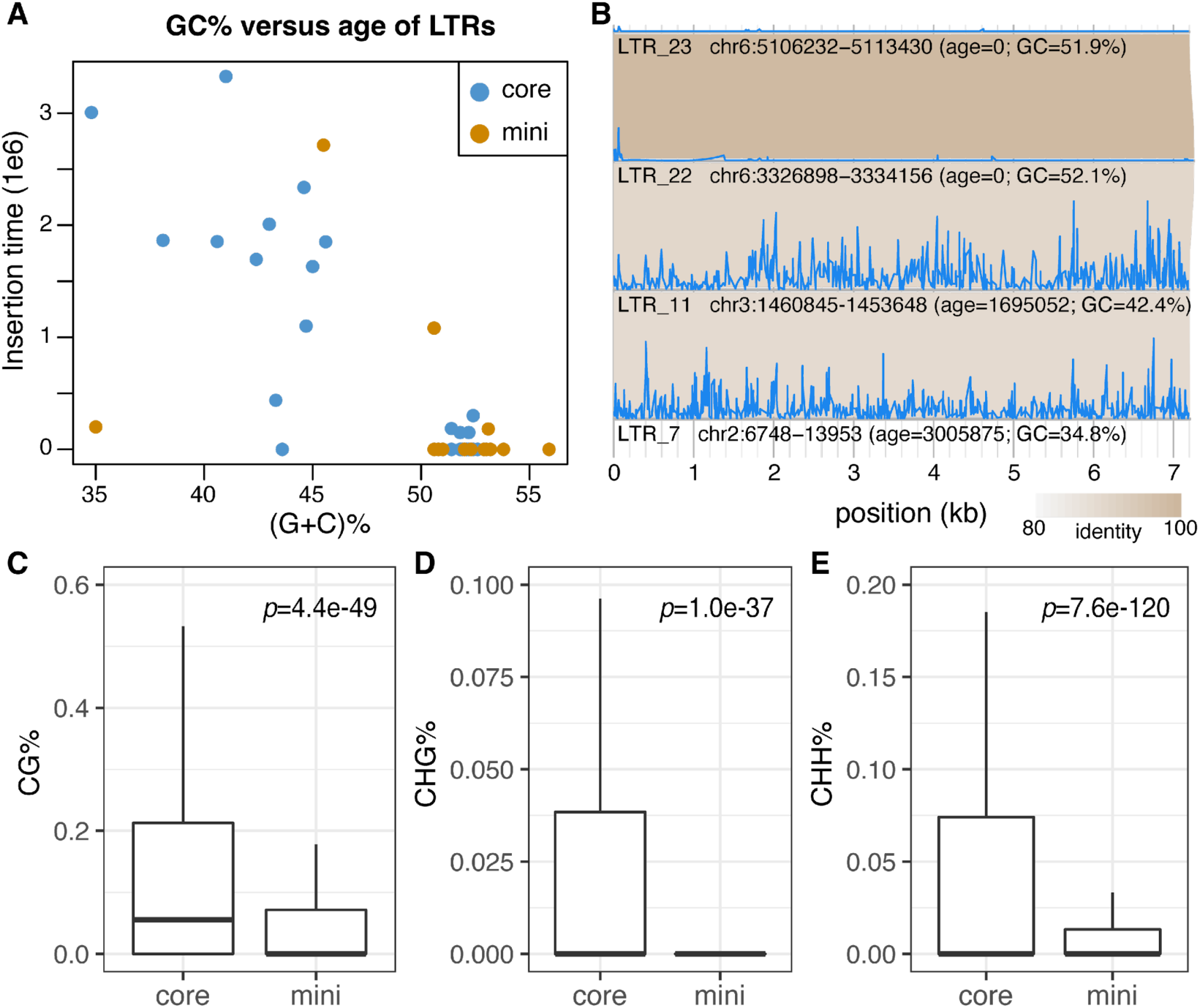
GC content, age, and DNA methylation of LTR retrotransposons in MC7. (**A**). Insertion time of each intact LTR retrotransposon was estimated and plotted versus the GC content. Each dot represents an intact LTR retrotransposon. Locations of retrotransposons in core-chromosomes (core) or mini-chromosomes (mini) are color-coded. (**B**). Alignments of four homologous *Copia* LTR retrotransposons and their CG methylation profiles. The identity of alignments were color coded. Blue curves indicate CG methylation profiles of LTR elements, showing LTR_22 and LTR_23 were not hypermethylated. (**C-E**). Cytosine DNA methylation between intact retrotransposons on core chromosomes and mini-chromosomes. P-values (*p*) were from t-tests between the two groups. Y-axes in the panels **C**, **D**, **E** represent the percentage of cytosine methylation in respective regions.

Genomic DNAs from two biological replicates of TF05-1 mycelium samples were subjected to whole genome bisulfite sequencing (WGBS) for detecting cytosine methylation [24,28]. On average, 6.5%, 1.1%, and 2% were detected for cytosine bases in the context of CG, CHG, and CHH, respectively, where H represents A, C, or T (**Table S13**). Many old LTR retrotransposons were hypermethylated, whereas young LTR retrotransposons were largely hypomethylated in all three contexts (**Figure S7, S8, S9**). For example, four homologous *Copia* elements showing higher than 85% identity between each pair were categorized into two old and two young LTR elements. Both old LTR retrotransposons were methylated at a high level and both young ones were at a very low level (**Figure 6B**, **Figure S7, S8, S9**). Because most young LTR retrotransposons were located on the mini-chromosomes, cytosine bases of LTR retrotransposons on mini-chromosomes were significantly less methylated as compared with those of LTR counterparts on core chromosomes (**Figure 6C**, **6D**, **6F**). The finding that LTR retrotransposons with higher G+C content (i.e. >50%) are hypomethylated suggests these elements have not undergone repeat-induced point mutation (RIP), a genomic mechanism in many fungi that causes methylated cytosine to tyrosine transition mutations that can result in transposable element inactivation [29,30]The results show that most LTR retrotransposons on mini-chromosomes were young insertions with a high G+C content and low DNA methylation, indicating that LTR retrotransposons on mini-chromosomes have not undergone RIP and are more likely to remain active [31].

## Discussion

Here, we examined genomic variation through CHEF, FISH, and comparative genomics approaches using sequencing reads and genome assemblies. CHEF karyotyping of *M*. *oryzae* strains can readily detect mini-chromosomes if they are distinguishable from core chromosomes in size. It can also determine the number of mini-chromosomes in a genome in case that multiple mini-chromosomes have sufficiently distinct sizes. Cytological karyotyping, the microscopic study of chromosomes, is a vital tool for understanding the genomic architecture of plants, insects and other eukaryotes [23]. FISH visualizes chromosomes directly regardless of chromosome sizes and could detect interchromosomal translocation of large DNA fragments using proper probes. We showed that FISH can be applied to fungal karyotyping even with the small size of fungal chromosomes. Neither CHEF nor FISH provides DNA sequences that offer the single base pair resolution of genome variants. Previously, sequencing of CHEF bands was developed, uncovering the genomic content of individual chromosomes [16,18]. For the complexity of a genome similar to *M*. *oryzae*, telomere-to-telomere (T2T) genome assemblies could be achieved through using Nanopore long reads or PacBio HiFi reads [20,22,32,33]. From this study, we learned that multiple additional factors could interfere with generation of T2T assemblies, including hundred-kilobase near identical duplications, multiple mini-chromosomes containing highly identical large fragments, and a high level of genomic instability that results in a certain level of variants from an *in vitro* culture. To avoid misleading conclusions from genome assemblies, comparative genomics directly using reads, such as CGRD, complementarily provides evidence of structural variation [24].

Integrating results from CHEF, FISH, comparative genomics approaches with genome assemblies and directly using sequencing reads, we characterized the genomic dynamics in monoconidial strains derived from the MoL isolate TF05-1, revealing pronounced genomic structural variation in mini-chromosomes. Such variation occurred in a few generations of *in vitro* culture or even within the same generation. The high level of structural variation in mini-chromosomes is associated with repetitive sequences in mini-chromosomes. We documented an intrachromosomal rearrangement in MC7202, a variant strain derived from MC7, in which a large chromosomal fragment of mini1 was deleted while the rest of the chromosome was duplicated. The rearrangement was hypothesized to be mediated by intrastrand base pairing of a tandem inverted repeat [26], showing an example that repetitive sequences provide homology for ectopic recombination. In our CHEF result of MC7, we frequently observed a weak band with the same size as the duplicated chromosome, indicating that some cells were subjected to such genome rearrangement. The result implies that structural variants around repetitive sequences with certain genomic organization could occur frequently, resulting in recurrent mutant events. Our data show that mini-chromosomes contain many intact LTR retrotransposons that were recent insertions and largely unmethylated at cytosine, indicating they have not been targeted by RIP. Active transposable elements could facilitate rearrangements in mini-chromosomes and possibly core chromosomes as well. Our result also shows that mini-chromosomes contain a few old LTR retrotransposons, which contain lower G+C content, probably due to RIP, and a high level of DNA cytosine methylation. Although ages of LTR retrotransposons may not be accurately estimated owing to potential RIP, these retrotransposons were unlikely to be recent insertion events and should be inactive. Genome-wide inactivation of transposable elements through RIP usually occurs during sexual crosses [34]. A few cases of presumably inactive LTR retrotransposons in mini-chromosomes indicate sexual crosses did not occur recently. Increasing lines of evidence support that mini-chromosomes may interfere with sexual fertility, reducing the likelihood of sexual crosses [8,20]. Under this scenario, sexual crosses occurred before the introduction of mini-chromosomes in TF05-1, and the inactivated retrotransposons in mini-chromosomes could have originated from core chromosomes. If there is a mechanistic link by which sex is impeded by mini-chromosomes, then mini-chromosomes can protect transposons from RIP and preserve their activity in the genome, possibly facilitating genome dynamics and variability. An alternative hypothesis is that the mini-chromosome escapes RIP by avoiding components of RIP machinery such as methylation of specific histone residues or targeting by the DNA methyltransferase-like protein RIP Defective (RID) [29] or Dim2 [35,36] through an unknown mechanism. Genome wide analysis of DNA methylation in other fungi has also reported incomplete transposable element methylation in fungi that lack mini-chromosomes [37,38]. This indicates that avoiding RIP is a common occurrence in fungal genomes, and given the high occurrence of non-methylated LTR retrotransposons on mini-chromosomes, the mini may be particularly suited to escape genomic defense mechanisms. Future research to further understand the mechanistic basis of variable genome defense will be important to understand fungal genome evolution.

Our results imply selective transfer of a mini-chromosome between strains of two recently emerged *M*. *oryzae* populations: MoT causing wheat blast disease in South America, South Asia, and Africa, and MoL causing the turf and forage grass disease gray leaf spot in South America, the U.S., and beyond. The high levels of sequence similarity between mini2 in the U.S. MoL isolate TF05-1 and the single mini-chromosome in Bolivian MoT isolate B71 sharply contrast with the lower levels of sequence similarities between core chromosomes in these strains. This could have occurred either through horizontal transfer initiated by hyphal anastomosis and heterokaryon formation, or through recombination involving the sexual cycle. The recent emergence of both the MoL and MoT clades involved sexual recombination of standing variation in five previous host-adapted populations. Specifically, detailed genomic analyses indicate that both the MoL and MoT populations were relatively recently derived from a common progenitor lineage, known as chromosomal haplotype MoL1-1 and represented by the field isolate ATCC64557 recovered from *Lolium* spp. in the U.S. in 1972 [14]. Core chromosomes of this progenitor lineage strain contain mixed chromosomal recombination blocks characteristic of sexual recombination between an *Eleusine*-adapted strain and an *Urochloa* (*Brachiaria*)-adapted strain. The progenitor population further diversified through sexual crosses including isolates from three additional plant host genera, *Stenotaphrum* spp., *Luziola* spp., and an unknown population related to but distinct from *Oryza*/*Setaria* pathogens. The MoL and MoT populations now occur as 13 *Lolium*-adapted and 34 *Triticum*-adapted chromosomal haplotypes that retain progenitor *Eleusine*/*Urochloa* recombination breakpoints, and that now appear to be mainly reproducing asexually [14]. These findings are consistent with a model that this multi-hybrid swarm of sexual recombination occurred via contemporaneous growth of these host-adapted *M*. *oryzae* strains in hay or dried straw from infected forage pastures (*Urochloa* spp. and *Stenotaphrum* spp.) containing associated weeds (*Eleusine* spp. and *Luziola* spp.) in Brazil. New genotypes were subsequently dispersed geographically through transport of infected plant material and/or seeds. Despite extensive genomic analyses documenting the role of sexual recombination in the emerging MoL and MoT populations, TF05-1 remains the earliest known field isolate containing the B71-like mini-chromosome. Therefore, horizontal transfer originating through asexual hyphal fusion remains an option for dispersal of the B71-like mini-chromosome.

We report a high level of B71-like mini-chromosome instability within purified monoconidial cultures of the U.S. *Lolium*-adapted field isolate TF05-1 grown on agar medium. This *in vitro* study complements our recent analysis of high levels of mini-chromosome variability in MoT field isolates collected from a natural field population in the year of first identification of South American B71-like strains causing wheat blast in Zambia in 2018, and in the following two years [22]. Zambian MoT field isolates exhibited variation in mini-chromosome numbers and DNA sequence, compared to relative stability of the core chromosomes. The levels of instability in these B71-like mini-chromosomes stands in contrast to a different 1.2-Mb mini-chromosome, mChrA, which appears to be highly syntenic between a 1989 *Eleusine* isolate from Brazil (Br62) and a 2011 rice isolate from Italy [39]. Clearly, understanding the composition and behavior of the complex mini-chromosome compartment of the *Magnaporthe* pangenome is critical for predicting the evolutionary potential for this important family of cereal, forage, and turf crop pathogens. Horizontal transfer has been implicated both among host-adapted lineages of the species *M*. *oryzae*, as well as between different *Magnaporthe*/*Pyricularia* species. Specifically, potential cases of horizontal transfer have been identified by focusing on avirulence effector genes [17,40], on core chromosomal segments [33], and on the 1.2-Mb mini-chromosome mChrA from the Italian *Oryza* isolate AG006 [39]. Mini-chromosomes may play important roles for horizontal transfer during vegetative hyphal growth and heterokaryon formation. It is currently unknown if mini-chromosomes can transfer between separate haploid nuclei in heterokaryotic hyphal cells, or if formation of transient diploid nuclei is required during the parasexual cycle, which has been demonstrated in *M*. *oryzae* under laboratory conditions [41]. Mini-chromosomes might benefit from their small size in transmission during asexual crosses, akin to the transmission advantage resulting in meiotic drive of certain chromosomes during sexual crosses[42]. The ability to exchange genomic content asexually offers means to increase genetic diversity and adaptivity of the *M*. *oryzae* species [39]. Mini-chromosomes are considered to be a reservoir and a cradle for adaptive evolution of effector genes [16,18,43]. Therefore, mini-chromosomes could be important mediators for the distribution and the functional evolution of effector genes among fungal strains of the *Magnaporthe*/*Pyricularia* genus, which in turn would impact host specificity and pathogenic aggressiveness.

## Materials and Methods

### Mo strains

MoL field isolate TF05-1 was collected from *Lolium arundinaceum* (previously *Festuca arundinaceum*) in Lexington, Kentucky in the US in 2005 [44]. The strain was shared by Dr. Mark Farman at University of Kentucky. Single spore cultures of TF05-1 were isolated to form multiple monoconidial strains (e.g., MC5 and MC7). Research with native US MoL strain TF05-1 was conducted in our normal Biosafety Level-1 (BSL-1) laboratory. This contrasts with all research conducted on exotic MoT strains, which remains restricted to BSL-3 laboratory containment conditions according to strict USDA-APHIS permit requirements.

### Whole genome sequencing using an Illumina platform

The procedure of fungal growth, DNA extraction for Illumina sequencing was adapted from the method described previously [22]. Briefly, TF05-1 isolates were cultured on oatmeal agar (OMA) plates and incubated under continuous light at 25°C for 6-7 days [45]. Pieces of OMA culture were used for liquid cultures. Mycelial mats were collected for DNA extraction with Qiagen plant DNA purification kit (Qiagen, Genmany). Paired-end sequencing was performed on the isolates using the Illumina platform at Novogene USA.

### Construction of phylogenetic trees

Publicly available whole genome sequencing (WGS) data of 20 Mo field isolates (**Table S1**) were used with the WGS data of TF05-1MC7 for discovering SNPs. Briefly, WGS reads were aligned to the B71 reference genome (B71Ref2) via BWA (0.7.17-r1188) [16]. Valid alignments requiring at least 95% identity and 95% coverage were passed for SNP calling through GATK [46]. As isolate B71 was in the 20 Mo isolates and the B71 genome served as the reference genome, we required that the B71 genotypes of all SNPs matched the reference genome. Furthermore, VCFtools with the parameters “--maf 0.1 --max-missing 0.2” was employed to filter SNPs [47]. Phylogenetic trees were generated using IQ-TREE 2 [48]. Two separated trees: one included SNPs in core chromosomes and the other included SNPs in mini-chromosomes. Both trees were constructed using the same optimized model, ‘TVM+F+ASC+R3’. The resulting trees were visualized using ITOL [49].

### Analysis of copy number variation via CGRD

Copy number variation was analyzed using the pipeline CGRD (v0.3.6) with Illumina and Nanopore whole genome sequencing data [24].

### Pulsed-field electrophoresis using CHEF

Mycelia were cultured in a 500 mL Erlenmeyer flasks containing 250 mL CM (1% glucose, 0.6% yeast extract and 0.6% Tryptone) at 130 rpm, 25℃ for 48 h, and then mycelia were filtered using a sterile funnel with one layer of milk filter. Subsequently, the collected mycelium was transferred to a mortar and ground for approximately 10 min. The ground material was then transferred to a 500 mL Erlenmeyer flask containing 250 mL of CM, and incubated at 25°C and 160 rpm for 16 hours. Following this incubation, the mycelial culture was collected using a sterile funnel with a single layer of milk filter. Collected mycelium was washed with 200 mL of sterile double-distilled water (ddH2O), followed by 50 mL of 1 M sorbitol. The mycelium was dried using sterile paper towels, transferred to a 200 mL Erlenmeyer flask containing 40 mL of lysis buffer (10 mg/mL lysing enzyme in 1 M sorbitol), and incubated at 90 rpm and 28°C for 2.5 hours. Following the incubation, the lysing solution was filtered through a single layer of sterile Nytex into fresh 50 mL conical tubes and centrifuged at 4,500 rpm (3,204 g) for 10 minutes at 4 °C. The resulting pellet was washed with 10 mL of SE buffer (1 M sorbitol, 50 mM EDTA) and centrifuged again at 4,500 rpm (3,204 g) for an additional 10 min at 4°C. The pellet was resuspended in 100-300 mL of SE buffer (1 × 10⁹ protoplasts/mL). The protoplast suspension warmed to 37°C in a water bath. Thirty-seven mL of preheated low-melting agarose and 63 mL of the protoplast suspension were added to a preheated (37°C) 2.0 mL microcentrifuge tube and mixed gently by pipetting. The suspension was transferred to a plug mold and allowed to solidify at 4°C for 30-60 minutes. Once solidified, the agarose plug was transferred into a 10 mL conical centrifuge tube containing Proteinase K buffer (2.5 mL Proteinase K Reaction Buffer per 1 mL of agarose plugs), followed by addition of 100 μL of Proteinase K stock. The plugs were incubated overnight at 50°C in a water bath without agitation. On the following day, the plugs were washed four times in 1x Wash Buffer for 1 h each at room temperature (RT) with gentle agitation (using 1 mL of 1x Wash Buffer per plug). The washed plugs were stored at 4°C in a 0.5× TBE buffer. Seven% Certified Megabase Agarose in 0.5×TBE was used for separation of mini-chromosomes. The CHEF electrophoresis was carried out in a Bio-Rad Mapper XA system with switching intervals of 1,200-4,800 seconds for 13 days (1.2 V, 120°, and 5 °C).

### Sequencing of DNA recovered from CHEF bands

The genomic DNA band from the CHEF gel was excised and placed inside a D-Tube Dialyzer (Millipore, CAT#71508-3, USA). The D-Tube was filled with a 0.5×TBE buffer and placed in the middle of the chamber of the Mapper XA system subjected to switching intervals of 1,200-3,600 seconds for 24 hours. After the run, DNA fragments were gently washed off the membrane in one side of the D-Tube using a pipette. The 0.5×TBE buffer containing DNA was then transferred to a new tube. Next, DNA in solution was loaded onto a 0.1 μm MCE membrane filter (MF-Millipore, CAT#VCWP04700, USA), which floated over a 40% PEG8000 solution to concentrate and wash DNA. DNA was washed twice with DNase-free water. Finally, the DNA was eluted with 50 μL of DNase-free water. The purified DNA sample was subsequently used for Illumina sequencing.

### Extraction of genomic DNA of monoconidial strains for Nanopore sequencing

Mycelium samples were frozen in liquid nitrogen and ground. Approximately 0.1 g sample powder was prepared in a fresh 2.0 mL tube followed by adding 550 mL extraction buffer I (100 mM Tris pH8.0, 250 mM NaCl, and 100 mM EDTA in ddH_2_O). 50 mL 20% SDS was then added to each tube and gently inverted to mix. The mixture was incubated at 37℃ for 1 hour in a water bath. Mix the sample gently each 20 min during incubation. Add 75 mL 5 M NaCl, invert the tube to mix. Add 65 mL (10% CTAB and 0.75 M NaCl; preheated at 65 ℃), invert to mix, and incubate at 65 ℃ for 20 min. Add 700 mL Tris-Saturated Phenol-Chloroform (V:V=1:1) to each sample and vortex for 5-10 min. Centrifuge at 12,000 rpm, RT for 12 min. Transfer the supernatant to a fresh 2.0 mL tube. Add 2 volumes of 100% ethanol to each sample, invert gently to mix. Put the samples in −20℃ for more than 1 h or overnight. Centrifuge at 12,000 rpm, RT, for 2 min. Add 500 mL Phenol:Chloroform Isoamyl Alcohol to each sample and vortex for 5-10 min. Centrifuge at 12,000 rpm, RT, for 10 min. Transfer the supernatant to a fresh 2.0 mL tube. Add 2 volumes of 100% ethanol to each sample and invert gently to mix. Put the samples in −20 ℃ for 1 h. Centrifuge at 12,000 rpm, RT, for 10 min. Discard the supernatant and wash the pellet with 1 mL 70% ethanol for two times. Remove the supernatant and air dry for 5-10 min. Add 100 mL ddH_2_O or TE to each sample.

### MC7 whole genome sequencing and genome assembly

Nanopore long read sequencing and genome assembly methods were previously described [22]. Briefly, genomic DNA was loaded to a BluePippin cassette provided in the kit BLF7510 using the 20 kb size selection High-Pass Protocol (Sage Science, USA). Subsequently, library preparation was carried out using the SQK-LSK109 kit (Oxford Nanopore, UK). The resulting library was loaded onto the FLO-MIN106 flow cell (Oxford Nanopore, UK) and sequenced using the MinION Mk1B sequencing device (Oxford Nanopore, UK). Guppy version 4.5.2 (Oxford Nanopore, UK) was employed to convert Nanopore raw data (fast5) into FASTQ format, using default parameters. Long reads were input for draft genome assembly using Canu v2.1.1 [50] with the following parameters: genomeSize = 46m; minReadLength = 5000; minOverlapLength = 1000; corOutCoverage = 60; correctedErrorRate=0.1. The orientation of the resulting assembly was redirected using a custom script based on the orientation of B71 reference genome [16], and fragmented contigs from mini-chromosomes were manually scaffolded based on the output suffixed with “best.edges.gfa” generated by the Canu assembler. The “best.edges.gfa” assembly graph data was visualized through Bandage (v0.8.1) [51]. The reorganized assembly was polished using Nanopolish [52] with Nanopore raw FAST5 data and Pilon [53] with Illumina short reads sequence. Note that the resulting polished genome of MC7 was referred to as TF05-1MC7Ref1.

### Genome annotation of TF05-1MC7Ref1

Mycelium samples from MC7, MC7202, and MC7212, were subjected to RNA extraction using the RNeasy Plant Mini Kit (Qiagen, USA) for RNA-seq at Novogene (USA). All RNA-seq data was used for gene annotation of the genome assembly of MC7 with Funannotate (v.1.8.8) (github.com/nextgenusfs/funannotate). The repetitive sequence database used for the repeat mask was generated using Extensive de novo TE Annotator (EDTA, v.2.0.0) [54]. Transcripts assembled with RNA-seq data from both *in vitro* culture and *in planta* infection of the B71 isolate were supplied as transcription evidence [16]. Evidenced protein data included MG8 protein annotation from rice isolate 70–15 [55], UNIPROTKB/SWISS-PROT protein database released in March 2021, and an effector collection (https://raw.githubusercontent.com/liu3zhenlab/collected_data/master/Magnaporthe/known.effectors.db01.fasta).

### Analysis of structural variation via Syri

NUCMER (v.4.0.0) was employed to align the genome sequences of B71Ref2 and TF05-1MC7Ref1 with the settings (--maxmatch -c 500 -b 500 -l 20) [56]. Subsequently, alignments were filtered through NUCMER’s ‘delta-filter’ tool with parameters set to “-m - i 90 -l 500”. The filtered alignments were subjected to structural variation analysis using SYRI (v.1.5) [25]. The analysis was implemented using the pipeline of chrcomp (https://github.com/liu3zhenlab/chrcomp).

### Analysis of *BAS1*-*PWL2* segments

Package Homotools was used for identifying the homologous segments and plotting (Liu et al. 2024).

### Screening of MC5 and MC7 variants

Single spores of the original MC5 or MC7 strains were isolated from oatmeal plates. The single-spore isolates were subcultured separately for up to four generations. Each generation consisted of inoculation of the center of an oatmeal agar plate (100 mm diameter), mycelial growth to fill the plate and transfer of mycelial plugs from the plate edges for growth on a new oatmeal agar plate (**Figure 4A**). For each generation, 40 single spores from both MC5 and MC7 were isolated and cultured. CHEF analysis was performed to screen the variants with different number and/or size of mini-chromosomes as compared with the MC5 or MC7 parental strain.

### Fluorescence *in situ* hybridization (FISH)

Chromosome preparation for FISH was carried out using the germ tube burst method as described by Ayukawa and Taga (2022) with modifications. Fungal spores from rice polish medium or oatmeal medium plates were initially washed with 1mL PDB, 24g/L Potato Dextrose Broth (Fisher Scientific). Subsequently, spores were collected by centrifuging at 3,000 g for 5 min at the RT. The collected spores were then resuspended in 500–1,000 uL PDB to achieve a spore concentration of 4.5 × 10^5^ spores/mL. About 200 μL of this spore suspension was added to each glass slide for germination at 25°C for 6-7 hours. After this incubation, the PDB solution was gently removed, followed by addition of 200 μL of PDB containing 50 mg/mL Thiabendazole (TBZ). The slides were then incubated at 25°C for 2 h and 20 min. This process was performed within approximately 5 minutes from the first slide to the last slide. After incubation, the slides were gently dipped once in ddH2O and flicked 5-10 times to remove any remaining PDB. The slides were then immersed in a fixative solution composed of methanol and glacial acetic acid (17:3 v/v) for 30 minutes. The slides were briefly flamed once and quickly passed above the flame about 3 times to dry. All the preparations were stored at −80°C until use.

Primers to amplify DNA probes developed to identify individual core or mini-chromosomes in this study were provided in **Table S7**. PCR products were cloned into the pGEM-T-Easy vector (Promega). The design process for these probes began with extracting the conserved sequences from core chromosomes across multiple genomes of wheat isolate B71 [16], rice laboratory strain 70-15 [55], and a *Lolium* isolate LpKY97 [57]. Subsequently, 100 kb sequences from conserved regions of core chromosomes were extracted and aligned to the TF05-1MC7Ref1 genome assembly. The regions that aligned to multiple locations in the genome were removed, resulting in the final design of seven probes targeting core chromosomes. Mini-chromosome probes were designed to target the unique regions on mini-chromosomes, ensuring they could not cross-hybridize with core chromosomes. This was achieved through self-alignment of the TF05-1MC7Ref1 genome using BLAST+. To develop the mini-chromosome painting probe, mini-chromosome DNA was recovered from a CHEF gel using the Qiagen DNA elution kit. The eluted DNA was amplified using a REPLI-g kit (Qiagen) and used as the painting probe.

Probe DNA was labeled with either digoxigenin-11-dUTP or biotin-16-dUTP (Roche) using a nick translation reaction. FISH hybridization followed a previously published protocol used for maize chromosomes [58]. Probes were detected with Alexa Fluor 488 streptavidin (Invitrogen) for biotin-labeled probes and rhodamine-conjugated anti-digoxigenin for dig-labeled probes (Roche). Chromosomes were counterstained with DAPI in Vectashield antifade solution (Vector Laboratories). Images were captured using a Zeiss Axioplan 2 microscope equipped with a CoolSNAP HQ2 cooled CCD camera and AxioVision 4.8 software. Final image contrast was adjusted using Adobe Photoshop CS5.

### DNA methylation analysis

Genomic DNAs extracted from two biological replicates of TF05-1 mycelia were subjected to whole genome bisulfite sequencing (WGBS) on a Novaseq 6000 system at Novogene (USA). The data processing for bisulfite sequencing and DNA methylation analysis was conducted using the Bismark pipeline (v0.22.1) [24,28]. Initially, raw reads were trimmed with Trimmomatic (v0.38) [59] to eliminate adapter sequences and low-quality reads. Bowtie2 (v2.3.5.1) [60], integrated within Bismark, was employed for read alignment, with duplicate alignments removed prior to methylation assessment. Methylation levels were then determined for each cytosine site in the CG, CHG, and CHH contexts.

## Acknowledgements

We thank funding provided by USDA NIFA award (2021-67013-35724) to S. Liu, B. Valent, D. Cook, and D. Koo; USDA NIFA award (2021-68013-33719) to B. Valent; the NSF awards (2311738) to S. Liu; and NSF award (2011500) to S. Liu, B. Valent, D. Cook, and D. Koo. This is contribution no. 25-098-J from the Kansas Agricultural Experiment Station, Manhattan, Kansas.

## Data availability

Nanopore and Illumina genomic sequencing data have been deposited in the Sequence Read Archive (SRA) database under accessions PRJNA750760. The TF05MC7 genome assembly is available from Genbank accessions: GCA_036172685.1. Representative scripts used in this study have been deposited to GitHub (https://github.com/PlantG3/TF05.git).

## Author contribution

GL, HZ, DK, BV, and SL conceptualized experiments; GL, HZ, DK conducted experiments; GL, HZ, DK, and SL analyzed data; ZW and DC involved in supervision; all authors reviewed and revised the manuscript.

