## Supplementary Figures for "Highly dynamic supernumerary mini-chromosomes in a *Magnaporthe oryzae* strain"

Lin et al.

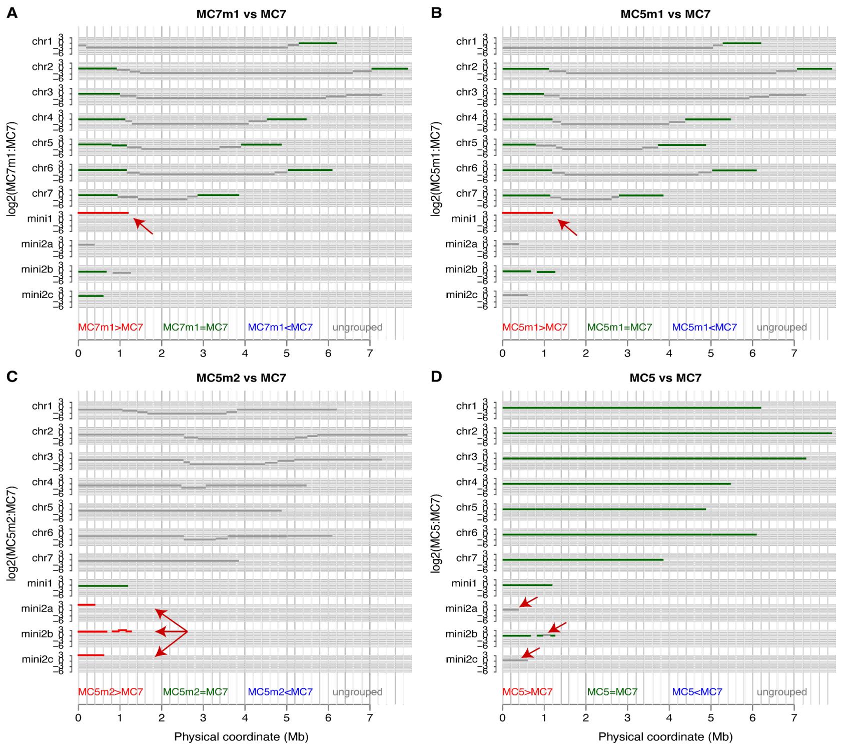


**Figure S1. Sequence analysis of mini-chromosomes on the MC7 genome assembly**
CHEF electrophoresis was conducted to visualize mini-chromosomes. DNA recovered from visible mini-chromosomal bands in CHEF gels was sequenced with an Illumina platform. While contaminating DNA fragments from any chromosomes could be sequenced, sequencing reads from the chromosomes from the excised band were expected to be markedly enriched in sequencing reads. CGRD was adapted to examine the enrichment in gel-recovered DNA as compared to whole genome sequencing (WGS) data, referred to as GelDNA-seq. GelDNA-seq can identify assembled contigs corresponding with visible bands on CHEF gels. (**A**). Sequence reads of the small mini-chromosome of MC7 (MC7m1) were compared with WGS reads of MC7. (**B**). Sequence reads of the small mini-chromosome of MC5 (MC5m1) were compared with WGS reads of MC7. (**C**). Sequence reads of the large mini-chromosome of MC5 (MC5m2) were compared with WGS reads of MC7. Red arrows in **A**, **B**, **C** indicate contigs with a marked enrichment. (**D**). WGS reads of MC5 (MC5) were compared with WGS reads of MC7, identified regions with a slightly higher DNA amount in MC5 as compared with MC7. The regions are highlighted by red arrows.

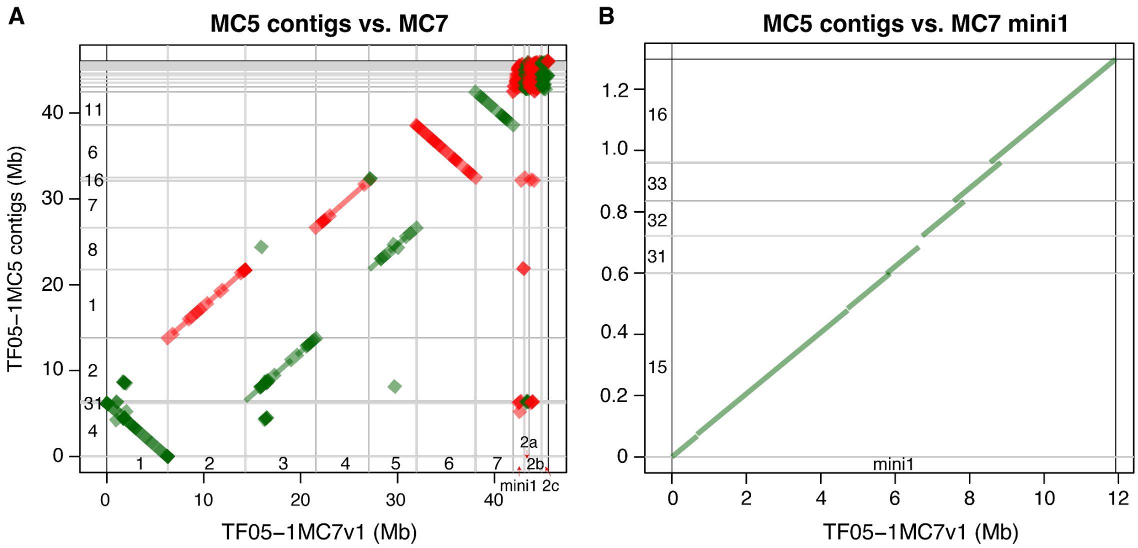


**Figure S2. Dotplots of MC5 contigs versus MC7v1**

(**A**) The dotplot to show alignments between TF05-1MC7v1 and MC5v1 (**B**) the dotplot to the assembled mini1 of TF05-1MC7v1 and a set of mini1-related contigs in TF05-1MC5v1. At least 10 kb and 90% identity were required for each alignment.

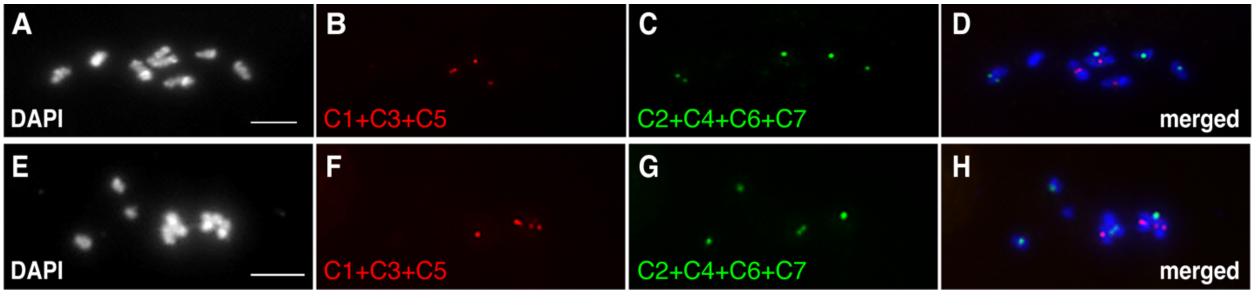


**Figure S3. FISH of MC7 with probes from core chromosomes**

Identification of individual core chromosomes by FISH using chromosome-specific DNA fragments as probes. Seven individual core chromosome-specific probes were pooled into two sets, each containing 3 or 4 probes. The first pool (C1+C3+C5) includes probe sets from chromosomes 1, 3, and 5, and the second pool (C2+C4+C6+C7) includes probe sets from chromosomes 2, 4, 6, and 7. One probe pool was used in each round of FISH, and after hybridization, the slides were washed and re-probed with the second pool. Note that no variation in the number of core chromosomes was observed in 20 nuclei from both MC5 and MC7. Bars=5 µm.

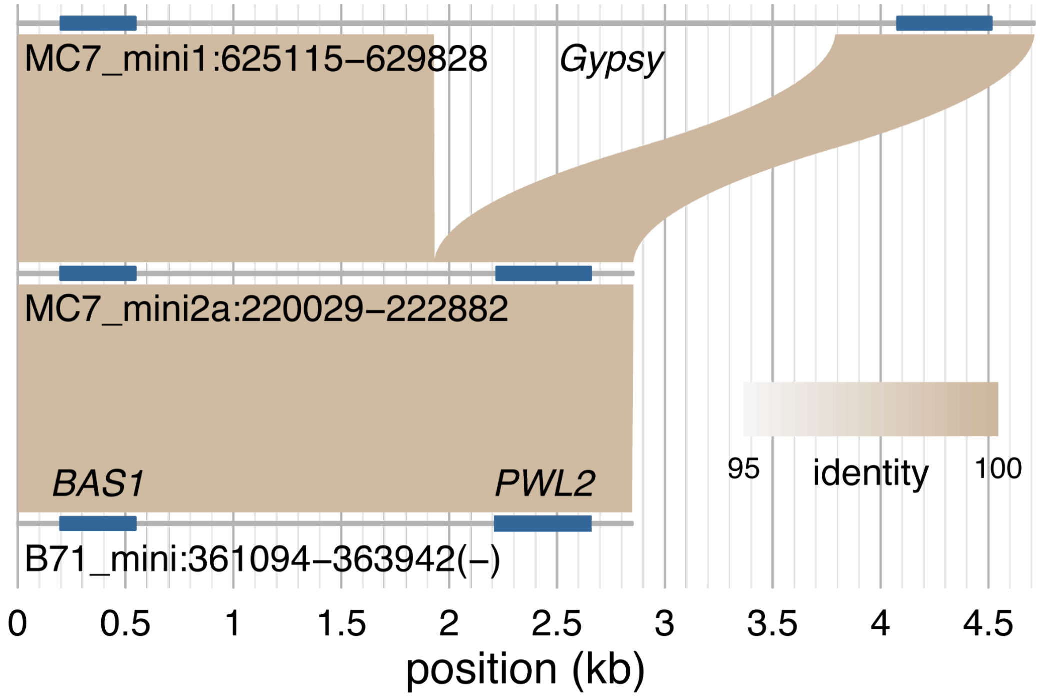


**Figure S4. Alignments of *BAS1-PWL2* segments**

Orange bars highlighted the locations of *BAS1* and *PWL2*. The alignment gap on mini1 was annotated as a *Gypsy* retrotransposon insertion.

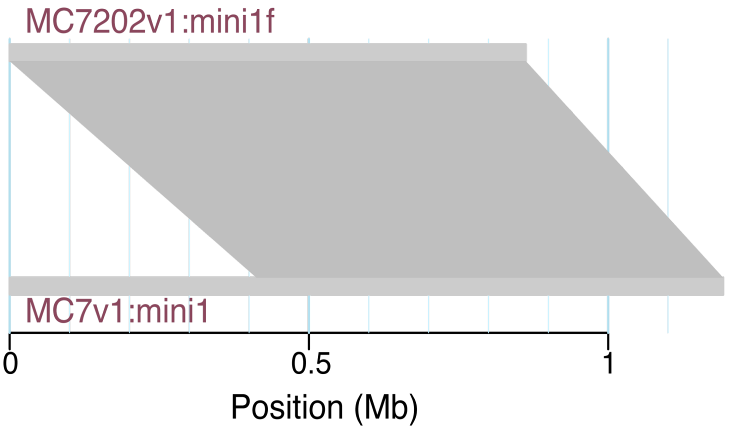


**Figure S5. Syntenic block alignments between MC7202 mini1f and MC7 mini1**
Mini1f assembled in MC7202 was compared with mini1 in MC7. Collective results indicate that the unaligned region on mini1 was absent in MC7202, and the aligned region was duplicated.

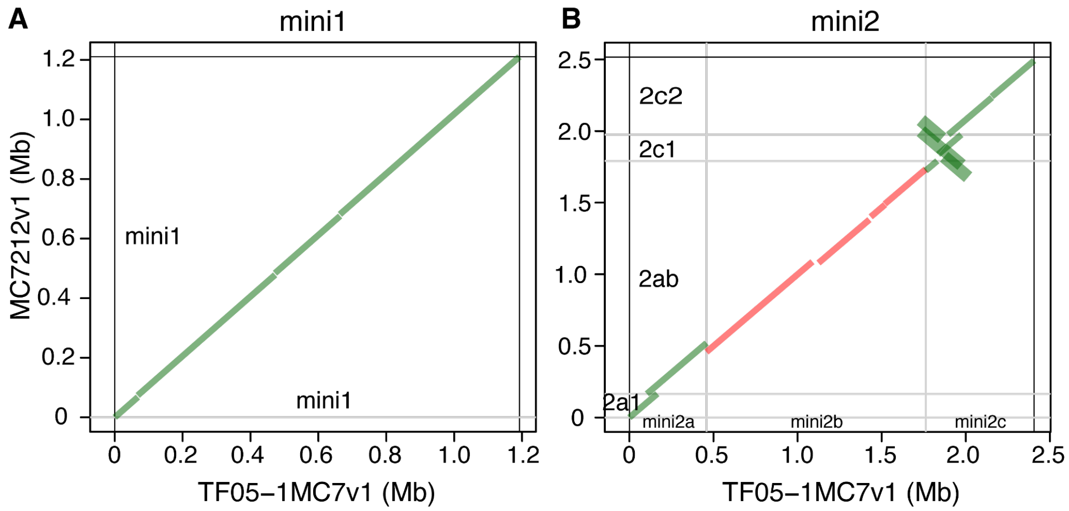


**Figure S6. Dotplots of MC7212 versus MC7v1 mini-chromosomes**
Dotplots were plotted to show alignments between the assembled mini1 of TF05-1MC7v1 and the assembled mini1 of MC7212v1 (**A**), and the assembled mini2 contigs of TF05-1MC7v1 and the assembled mini2 contigs of MC7212v1 (**B**). At least 10 kb and 90% identity were required for each alignment.

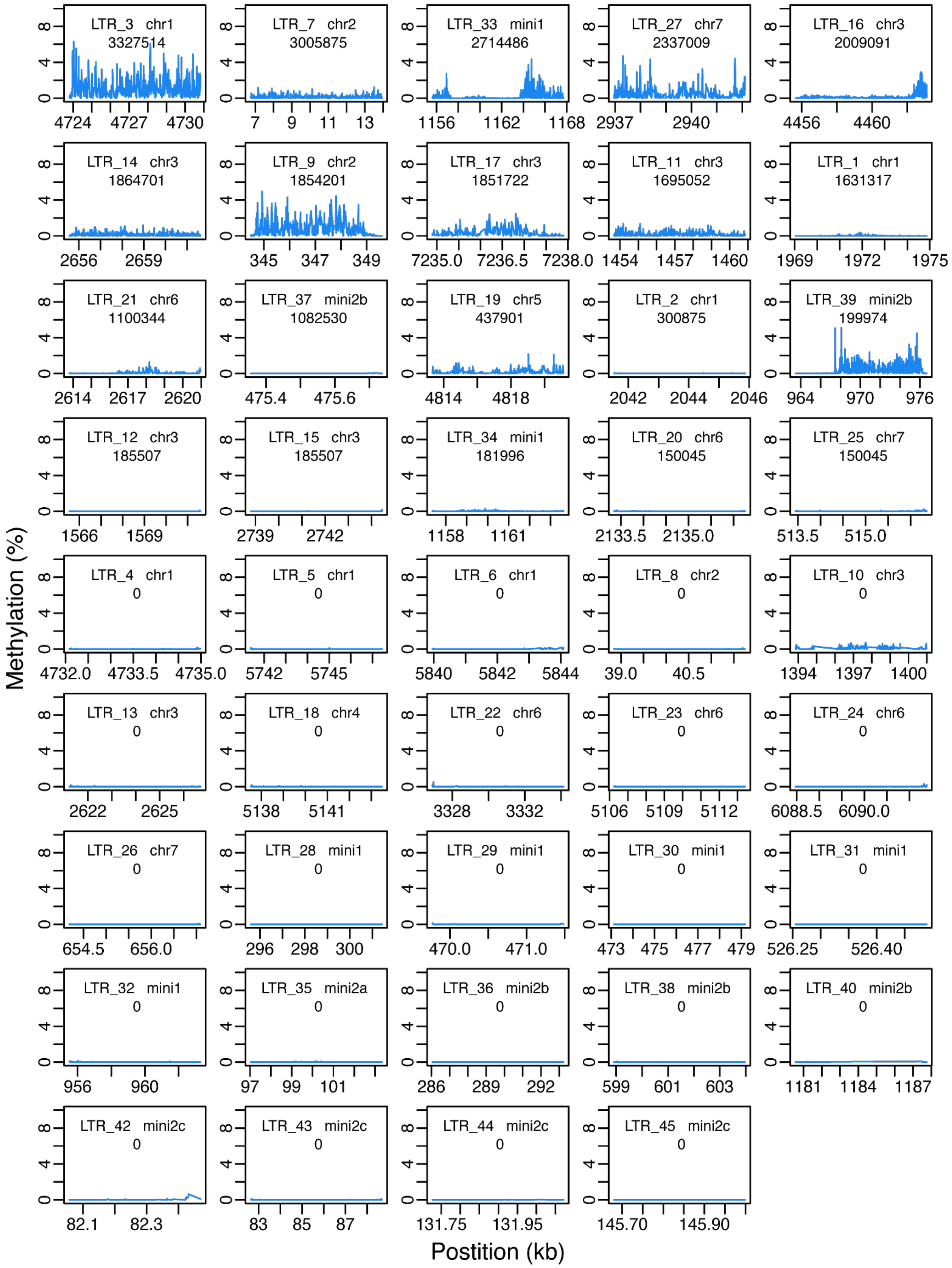


**Figure S7. CG methylation profiles of intact LTR retrotransposons**

The chromosomal source and the estimated age are labeled for each LTR. Y-axis signifies the level of cytosine methylation at CG sites.

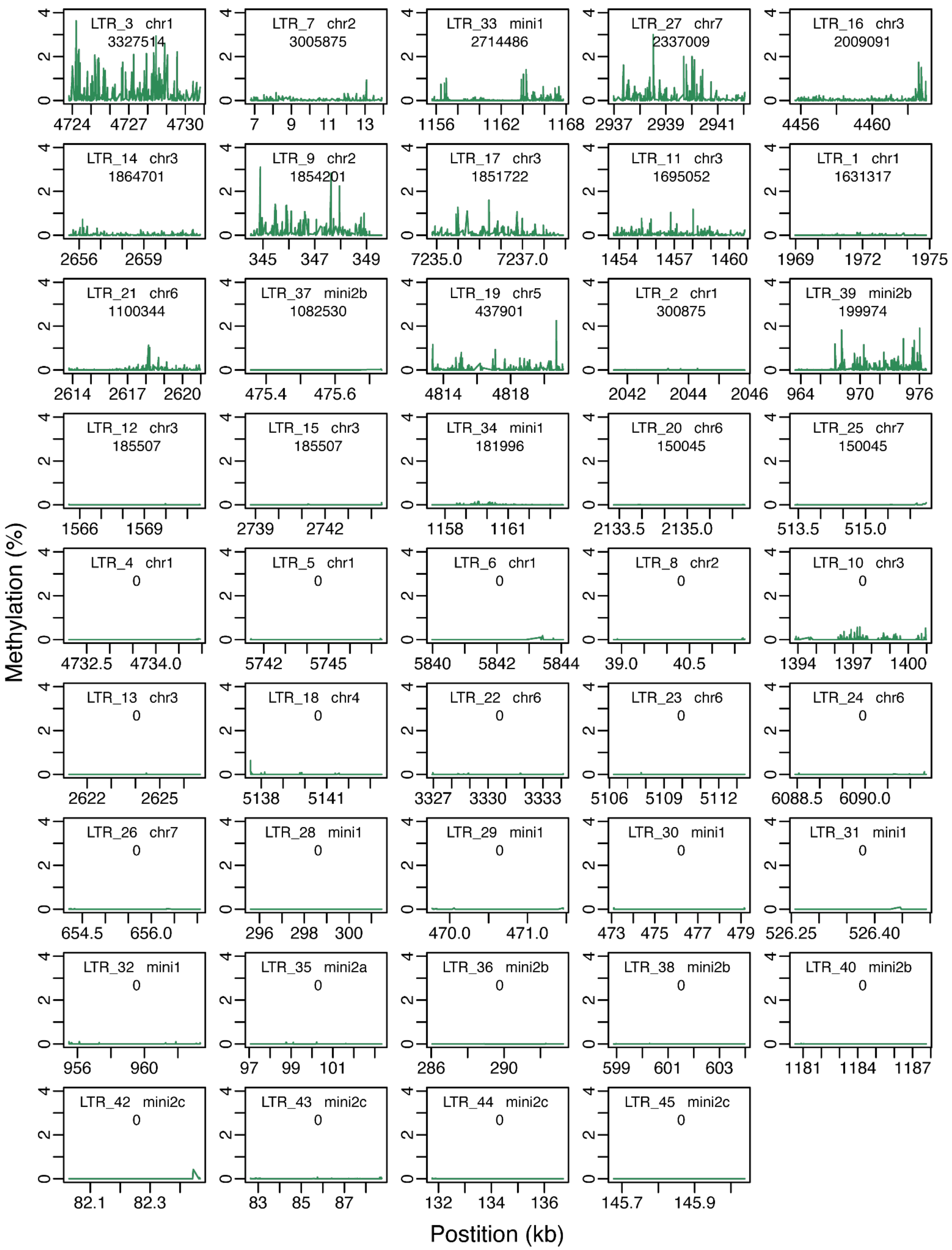

**Figure S8. CHG methylation profiles of intact LTR retrotransposons.**
The chromosomal source and the estimated age are labeled for each LRT. Y-axis signifies the level of cytosine methylation at CHG sites.

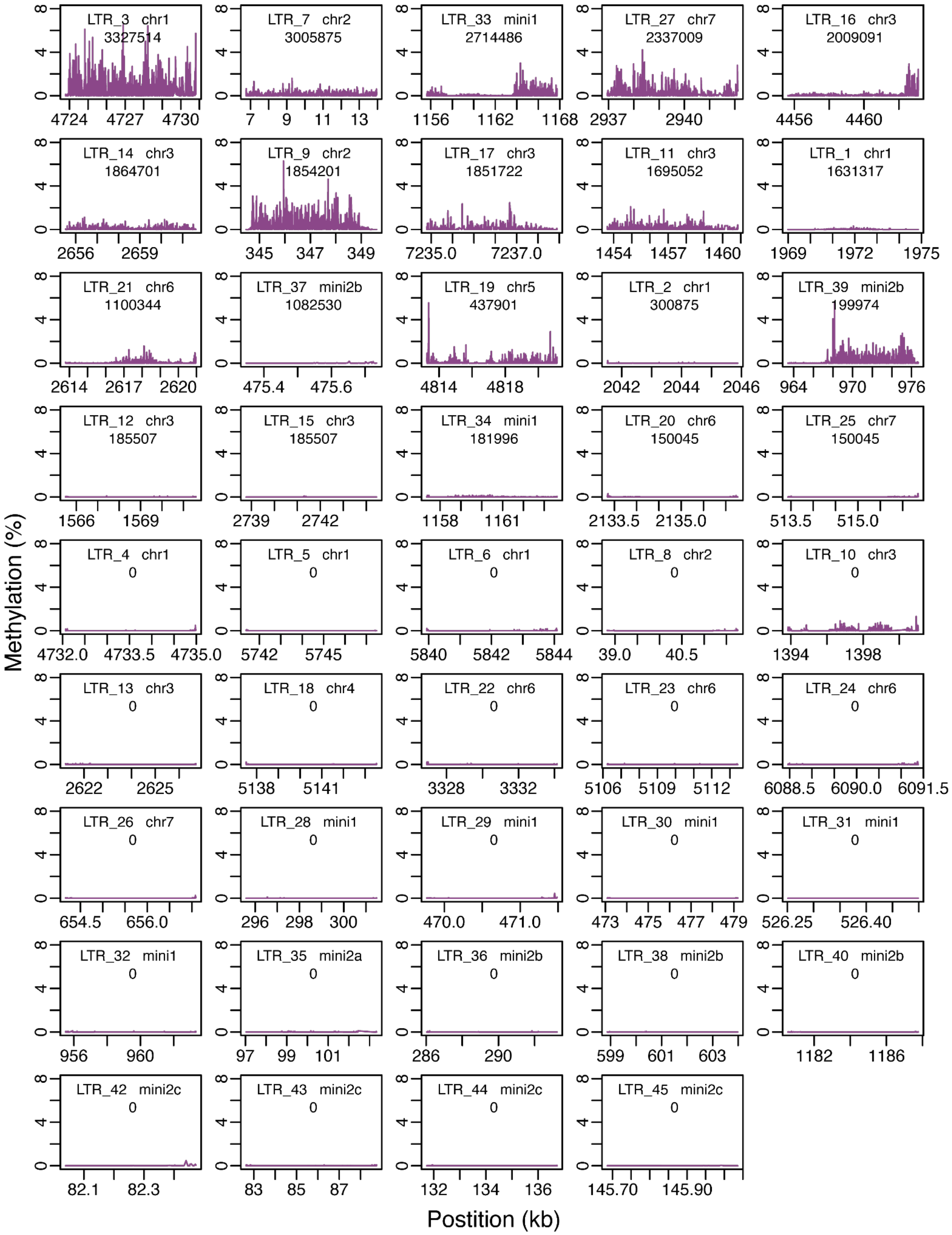


**Figure S9. CHH methylation profiles of intact LTR retrotransposons.**
The chromosomal source and the estimated age are labeled for each LRT. Y-axis signifies the level of cytosine methylation at CHG sites.
